# A novel method for in vivo assessment of soft tissue weight in bivalves

**DOI:** 10.1101/2024.12.25.630339

**Authors:** Shiyun Duan, Yunlong Gao, Xiaoli Hu, Xiaoshen Yin

**Author notes:** Corresponding author: Xiaoshen Yin; MOE Key Laboratory of Marine Genetics and Breeding, College of Marine Life Sciences, Ocean University of China, Qingdao 266003.

## Abstract

Aquaculture has been playing an increasingly important role in supplying proteins. To satisfy the rapidly growing demands for quality aquatic foods, the efficiency of aquaculture production needs to be improved. One strategy to achieve more efficient aquaculture production is to produce high-yielding seeds through breeding programs, for which in vivo assessment of target traits is indispensable to prevent candidate parents from being sacrificed for measurement and to track dynamically changing yield-related traits. Farmed mollusks, most of which are bivalves, rank the second among all aquatic animals in aquaculture production. Bivalves with heavier soft tissue, the major edible part, are preferable, target species for breeding and farming. A prerequisite for obtaining such species is to make accurate but nondestructive assessments of soft tissue weight, which cannot be accomplished using traditional dissection- or imaging-based methods. To resolve this issue, we take advantage of the complex relationships among shell dimension- and weight-related traits to construct classical, shell dimension- or wet weight-based models and novel, shell surface area-based models for in vivo assessment of soft tissue weight in Zhikong scallop (*Chlamys farreri*) and yesso scallop (*Patinopecten yessoensis*). Overall, soft tissue weight estimated by wet weight-involving models are highly correlated with observed values, indicating wet weight as a key proxy for soft tissue weight. While some models incorporating wet weight tend to systematically over- or under-estimate soft tissue weight despite the strong association between actual and estimated values, the novel, shell surface area-based model, MLR-*log*_10_*S*_*shell*_-RR, generates accurate estimates free of systematic biases across populations of different ages for both species, suggesting it as a promising approach for in vivo assessment of soft tissue weight for scallops and other bivalves with bilaterally symmetrical shells.

## 1 Introduction

Aquaculture has been growing nearly exponentially over the last several decades to supply quality proteins for the expanding population around the world (FAO, 2024). The global aquaculture production of aquatic animals reached 94.4 million tonnes in live weight and exceeded capture fisheries for the first time in 2022 (FAO, 2024). Despite the rapid increase in total aquaculture production, the efficiency of aquaculture production needs to be further improved to satisfy the demands for wholesome, economically viable, and environmentally sustainable aquatic foods. One strategy to achieve more efficient aquaculture production is to produce superior, resistant, high-yielding seeds for farming through breeding programs, for which rapid and accurate in vivo assessment of target, commercial traits is indispensable for three reasons. First, to speed up the breeding progress, aquatic animals serving as candidate parents for the next generation cannot be sacrificed for measurement in most breeding programs and thus a nondestructive assessment is essential. Second, yield-related traits, such as growth, usually vary across life stages, an accurate, in vivo assessment of these traits through the entire life cycle is indispensable for conducting genotype-phenotype association analysis at a finer temporal scale to illuminate the genetic basis underlying phenotypic variation to inform strategies for genetic improvements. Third, dynamically tracking changes in target traits through the entire grow-out stage or production cycle can help monitor the status of organisms in culture, identify top candidates for farming at early life stages, adjust farming plans promptly, pinpoint the correct time point for harvesting, and select potential superior parents for future breeding to accelerate the progress of genetic improvements and enhance the efficiency of aquaculture production.

Among all farmed aquatic animals, mollusks, following finfish, rank the second in aquaculture production with a total production of 18.9 million tonnes (FAO, 2024). Farmed mollusks, such as oysters, clams, scallops, and mussels, are mostly bivalves, which account for over 86% of aquaculture production of mollusks (FAO, 2024). As soft tissue is the major edible part of bivalves, bivalves with a larger proportion of soft tissue are preferable, target species of high commercial value. A prerequisite for obtaining such species is to make accurate but nondestructive assessments of soft tissue weight. As soft tissue is enclosed in hard shells consisting of two hinged and sometimes tightly closed valves, bivalves must be dissected first before their soft tissue can be weighed. Such traditional methods are usually destructive, leaving it impossible to save candidate parents for bivalve breeding or dynamically track changes in soft tissue weight of bivalves across various temporal scales.

Advanced technologies that have been broadly applied in medical examination and treatment, such as magnetic resonance imaging (MRI) and X-ray, provide a possibility for in vivo evaluation of interior parts of bivalves (Brand et al., 2009; Michael Holliman et al., 2008; Pouvreau et al., 2006; Zhao et al., 2021). MRI was first used to noninvasively investigate the anatomy of the Pacific oyster (*Crassostrea gigas*), which allows for a depiction of various organs in soft tissue at a sufficient resolution and a demonstration of some anatomical structures in the Pacific oyster (Pouvreau et al., 2006). Later, MRI was applied in the Eastern elliptio (*Elliptio complanata*) to assess the structure, function, and integrity of its soft tissue (Michael Holliman et al., 2008) and in the Chilean scallop (*Argopecten purpuratus*) to study its internal anatomy (Brand et al., 2009). MRI can effectively identify broodstock sex and condition, track physiological and metabolic status, and measure some commercial traits of bivalves in a nondestructive way to instruct bivalve breeding and farming (Pouvreau et al., 2006). X-ray, another technology that can capture the interior parts of bivalves, was used to assess adductor muscle for Zhikong scallop (*Chlamys farreri*) and yesso scallop (*Patinopecten yessoensis*). The high correlation between adductor muscle area and weight (*r*: ∼0.9) and between the percentages of adductor muscle area and weight (*r*: ∼0.7) verify X-ray as a potentially effective method for evaluating adductor muscle weight and percentage in scallops (Zhao et al., 2021).

While imaging-based methods can investigate the interior parts of bivalves, they can hardly be used to evaluate soft tissue weight in a reliable way for three reasons. First, imaging-based approaches can only take two-dimensional pictures to estimate the weight of internal parts of bivalves based on their areas. However, this does not work for assessing soft tissue weight as soft tissue has an irregular shape and an inconstant density. Second, imaging-based evaluations are heavily dependent on automatic or manual image processing, which may introduce uncertainties and biases. Third, the relatively high cost and low efficiency of these imaging-based assessments may restrict their broad application in aquaculture production. To resolve these issues, an alternative approach to noninvasively estimating soft tissue weight for bivalves based on morphometric relationships can be adopted. Allometric growth and morphometric relationship have long been studied in bivalves, including oysters, mussels, clams, razor clams, cockles, and scallops, across various temporal and spatial scales, to inform natural resources management, invasive species control, and aquaculture production (see Table S1 for details). Shell dimensions (e.g., shell length, shell width, and shell height) turn out to be promising predictors for the weight of whole bivalves, shells, and soft tissue, and multiple models of shell dimension-based estimations for all types of weight, such as simple linear regression, power function, and polynomial models, have been constructed for bivalve biomass assessment (Cameron et al., 1979; Coughlan et al., 2021; Eklöf et al., 2017; Golightly and Kosinski, 1981; Isom, 1971; Larson et al., 2014; McKinney et al., 2004; Molina et al., 2005; Powell et al., 2016; Turra et al., 2018; Wang et al., 2014).

Scallops are ecologically and commercially important components in ecosystem conservation and aquaculture production, but their morphometric relationships have rarely been investigated despite the critical role they play. Here, we construct classical, shell dimension- or wet weight-based models and novel, shell surface area-based models for two predominant aquaculture species in China, Zhikong and yesso scallops, to estimate their soft tissue weight. In comparison to models based on only shell dimensions, all models incorporating wet weight as an independent variable can generate estimates that are highly correlated with observed soft tissue weight (*r* > 0.8), implying wet weight as a key proxy for soft tissue weight. Although some of the wet weight-involving models tend to systematically over- or under-estimate soft tissue weight, indicating that they may not work for cases in which an accurate, exact predicted value is needed, the novel, shell surface area-based model, MLR-*log*_10_*S*_*shell*_-RR, which accounts for collinearity among shell dimension- and weight-related traits using ridge regression, can generate accurate assessments free of systematic biases across populations of different ages for both species, suggesting it as a robust and reliable phenotyping method for in vivo assessment of soft tissue weight for bivalves with bilaterally symmetrical shells.

## 2 Material and methods

### 2.1 Data collection and model fitting

We collected a total of 1,125 Zhikong scallops from two populations and 268 yesso scallops from two populations for analyses, including (1) 627 two-year-old Zhikong scallops grown at Rongcheng, Shandong and collected in 2020 (hereafter *Cf1*), (2) 498 17-month-old Zhikong scallops grown at Rongcheng, Shandong and collected in 2022 (hereafter *Cf2*), (3) 110 two-year-old Haida golden scallops, a new variety of yesso scallops with orange adductor muscle, grown at Dalian, Liaoning and collected in 2019 (hereafter *Hg*), and (4) 158 two-year-old Zhangzidao red scallops, a new variety of yesso scallops with orange shells, grown at Dalian, Liaoning and collected in 2019 (hereafter *Zr*). All four populations under study consist of samples randomly collected from the corresponding natural or domesticated populations.

For both Zhikong and yesso scallops, wet weight (*W*_*wet*_), shell weight (*W*_*shell*_), and soft tissue weight (*W*_*soft*_) were measured using a digital balance; shell length (*SL*), shell width (*SW*), shell height (*SH*), and the width of each shell (*SW*_1_, *SW*_2_) were measured using a digital caliper (Fig. 1). The bottom part of the shell (indicated with a blue box in Fig. 1, hereafter shell bottom) was approximated as a trapezoid and the height (*H*_*sb*1_ for one shell; *H*_*sb*2_ for the other shell) and length of two bases (*L*_*sb*11_, *L*_*sb*21_ for one shell; *L*_*sb*12_, *L*_*sb*22_ for the other shell) of this trapezoid were measured for each shell using a digital caliper (Fig. 1). We first tested the normality of shell dimensions, including shell length, shell width, and shell height, and wet weight in Zhikong and yesso scallops using the Shapiro-Wilk test in R 4.3.0 (R Core Team, 2023). Next, we compared shell dimensions and wet weight between the two populations in Zhikong or yesso scallop using the Student’s t-test, as implemented in R 4.3.0 (R Core Team, 2023). Given that shell dimensions and wet weight are not normally distributed (see 3.1 below for details in normality tests), we also compared shell dimensions and wet weight between the two populations in Zhikong or yesso scallop using the Mann-Whitney U test, as implemented in R 4.3.0 (R Core Team, 2023).

**Figure 1.**
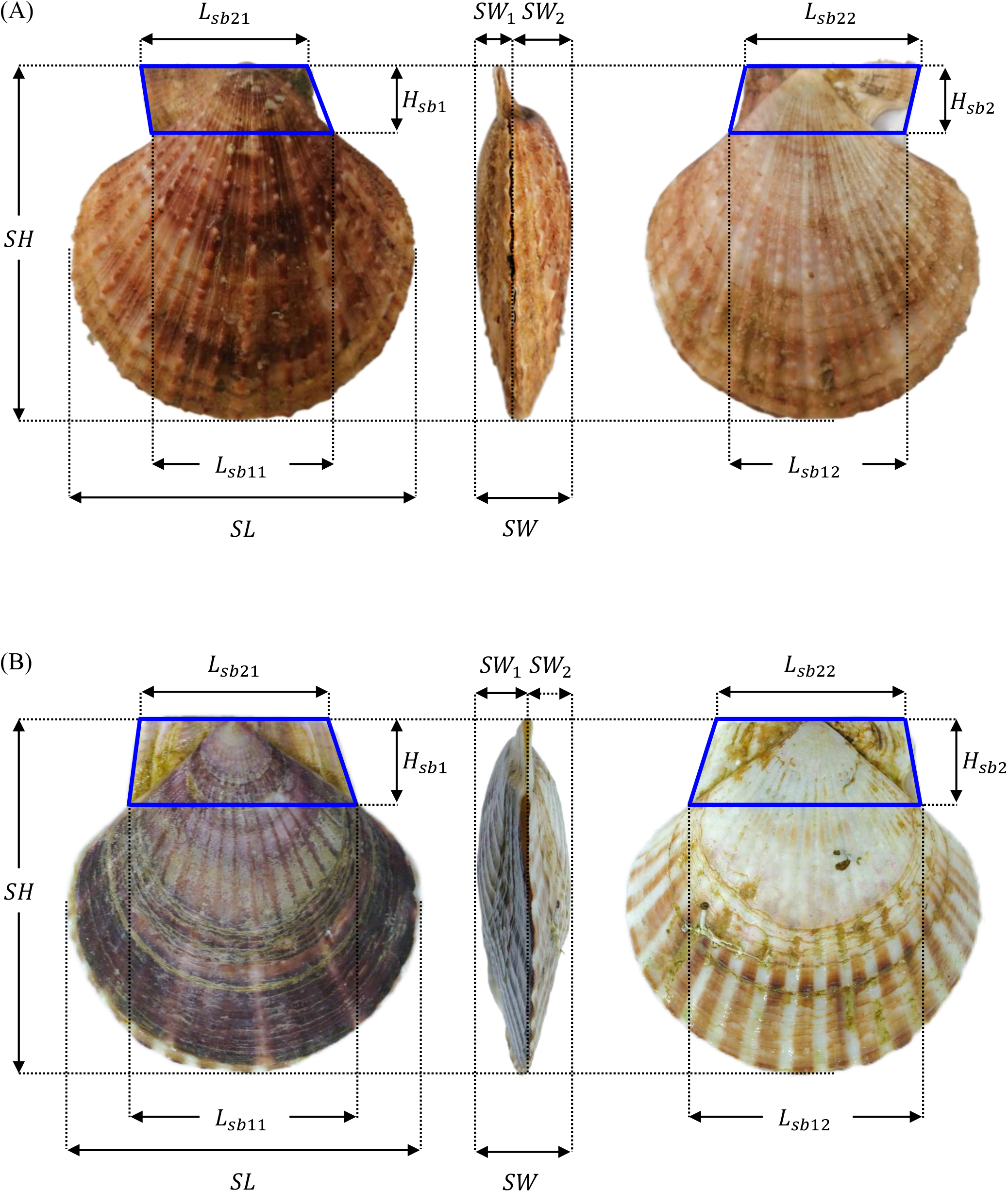
Shell dimensions of Zhikong (A) and yesso (B) scallops. *SL*: shell length; *SW*: shell width; *SH*: shell height; *SW*_1_ and *SW*_2_: the width of each shell; *H*_*sb*1_ and *H*_*sb*2_: the height of shell bottom of each shell; *L*_*sb*11_, *L*_*sb*21_, *L*_*sb*12_, and *L*_*sb*22_: the length of two bases of shell bottom of each shell.

To noninvasively assess soft tissue weight of Zhikong and yesso scallops, we fitted shell length, shell width, shell height, and wet weight as independent variables and soft tissue weight as a dependent variable to three models, simple linear regression, power function, and multiple linear regression. We conducted model fitting and soft tissue weight assessment in three ways: (1) for Zhikong scallop, we took each of the two populations, *Cf1* and *Cf2*, as a training set to construct an estimation model, with which we calculated soft tissue weight for the other population (i.e., testing set), (2) for yesso scallop, we took each of the two populations, *Hg* and *Zr*, as a training set to construct an estimation model, with which we calculated soft tissue weight for the other population (i.e., testing set), and (3) given that Zhikong scallop has larger population sizes, we randomly picked a subset of each population of Zhikong scallop (replicated three times per population), *Cf1* (n=50, 100, 150, 200, 250, 300) and *Cf2* (n=50, 100, 150, 200), as a training set to construct models for estimating soft tissue weight for the remaining subset of the same population and for the other population (i.e., testing set).

### 2.2 Shell dimension- or wet weight-based in vivo assessment of soft tissue weight

We first fitted shell length, shell width, shell height, or wet weight, as an independent variable, and soft tissue weight, as a dependent variable, of samples in the training set to simple linear regression (Equation 1-4) to determine coefficients, a and b, using ordinary least squares (OLS) regression, as implemented in R 4.3.0 (R Core Team, 2023). With coefficients estimated from the training set, we calculated soft tissue weight for testing sets using Equation 1-4, and named this simple linear regression-based estimation as SLR-*SL*, SLR-*SW*, SLR-*SH*, and SLR-*W*_*wet*_ methods for estimating soft tissue weight based on shell length, shell width, shell height, and wet weight, respectively.

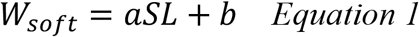

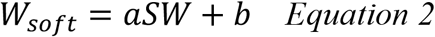

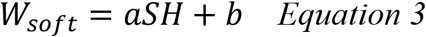

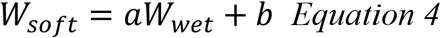

where a and b are constants.

We next fitted shell length, shell width, shell height, or wet weight as an independent variable, and soft tissue weight, as a dependent variable, of samples in the training set to power function (Equation 5-8) to determine coefficients, a and b, using OLS regression, as implemented in R 4.3.0 (R Core Team, 2023). With coefficients estimated from the training set, we calculated soft tissue weight for testing sets using Equation 5-8, and named this power function-based estimation as PF-*SL*, PF-*SW*, PF-*SH*, and PF-*W*_*wet*_ methods for estimating soft tissue weight based on shell length, shell width, shell height, and wet weight, respectively.

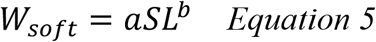

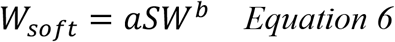

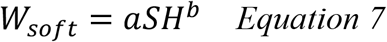

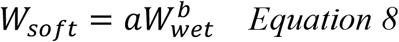

where a and b are constants.

Lastly, we fitted shell length, shell width, and shell height, as independent variables, and soft tissue weight, as a dependent variable, of samples in the training set to multiple linear regression (Equation 9) to obtain values of coefficients, a, b, c and d, using OLS regression, as implemented in R 4.3.0 (R Core Team, 2023), and ridge regression, which accounts for collinearity among shell dimension- and weight-related traits, as implemented in Scientific Platform Serving for Statistics Professional (hereafter SPSSPRO; https://www.spsspro.com/). With a, b, c, and d estimated from the training set, we calculated soft tissue weight for testing sets using Equation 9 and named this multiple linear regression-based estimation as MLR-*SLWH*-OLS and MLR-*SLWH*-RR methods for estimating soft tissue weight using OLS and ridge regression, respectively.

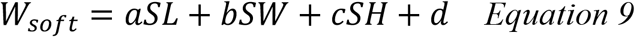

where a, b, c, and d are constants.

### 2.3 Shell surface area-based in vivo assessment of soft tissue weight

Soft tissue weight of bivalves can be calculated by subtracting shell weight from wet weight and shell weight can be calculated as the product of shell surface area (*S*_*shell*_), shell thickness (*th*_*shell*_), and shell density (*ρ*_*shell*_) (Equation 10). Assuming shell thickness and shell density are relatively constant in a population under the same environment, we fit wet weight and shell surface area to multiple linear regression (Equation 11) to estimate soft tissue weight.

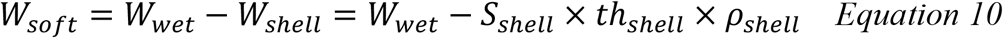

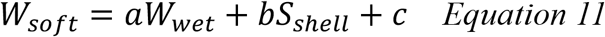

where a, b, and c are constants.

According to the shell morphology of Zhikong and yesso scallops, we divided a single shell into three parts, a quarter ellipsoid ABCDE, an irregular cone BCDEF, and two triangular shell ears FGH and FIJ (Fig. 2). The total surface area of the quarter ellipsoid ABCDE (*S*_*ellipsoid*_) and the two triangular shell ears FGH and FIJ (*S*_*shell ear*_) of two shells can be calculated using Equations 12 and 13, respectively (Fig. 2). To calculate the surface area of the irregular cone BCDEF, we first reconstructed a larger cone KCL(M) (hereafter source cone; see Appendix A for details) to which the irregular cone can fit and then expanded the lateral surface of source cone to a sector KLCM for the easiness of calculation (Fig. 2). The total surface area of the irregular cone of two shells can be calculated by subtracting the area of triangle KFB and triangle KFD from that of sector KBCD for both shells using Equation 14 (Appendix A; Fig. 2). Shell surface area of a scallop can be calculated as the sum of surface areas of the quarter ellipsoid (Equation 12), the two triangular shell ears (Equation 13), and the irregular cone (Appendix A; Equation 14).

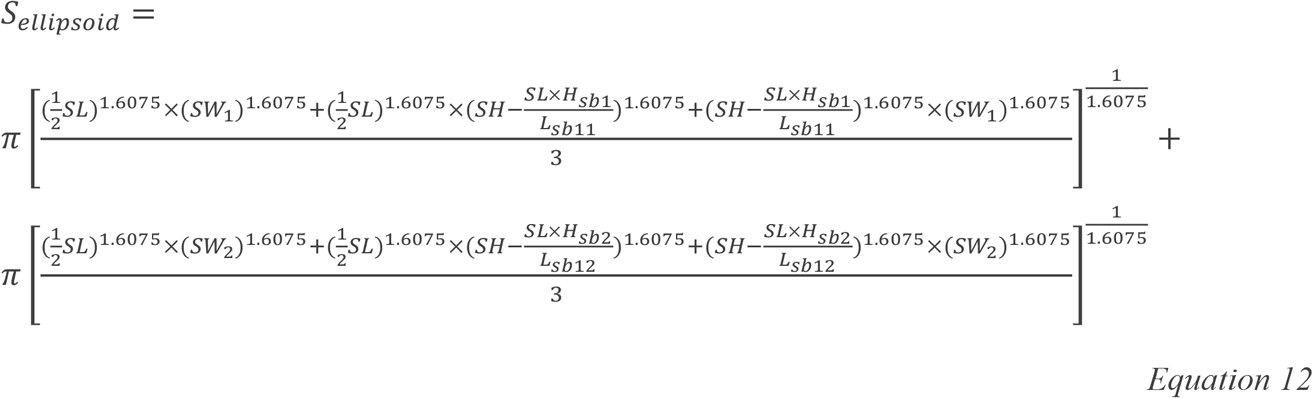

**Figure 2.**
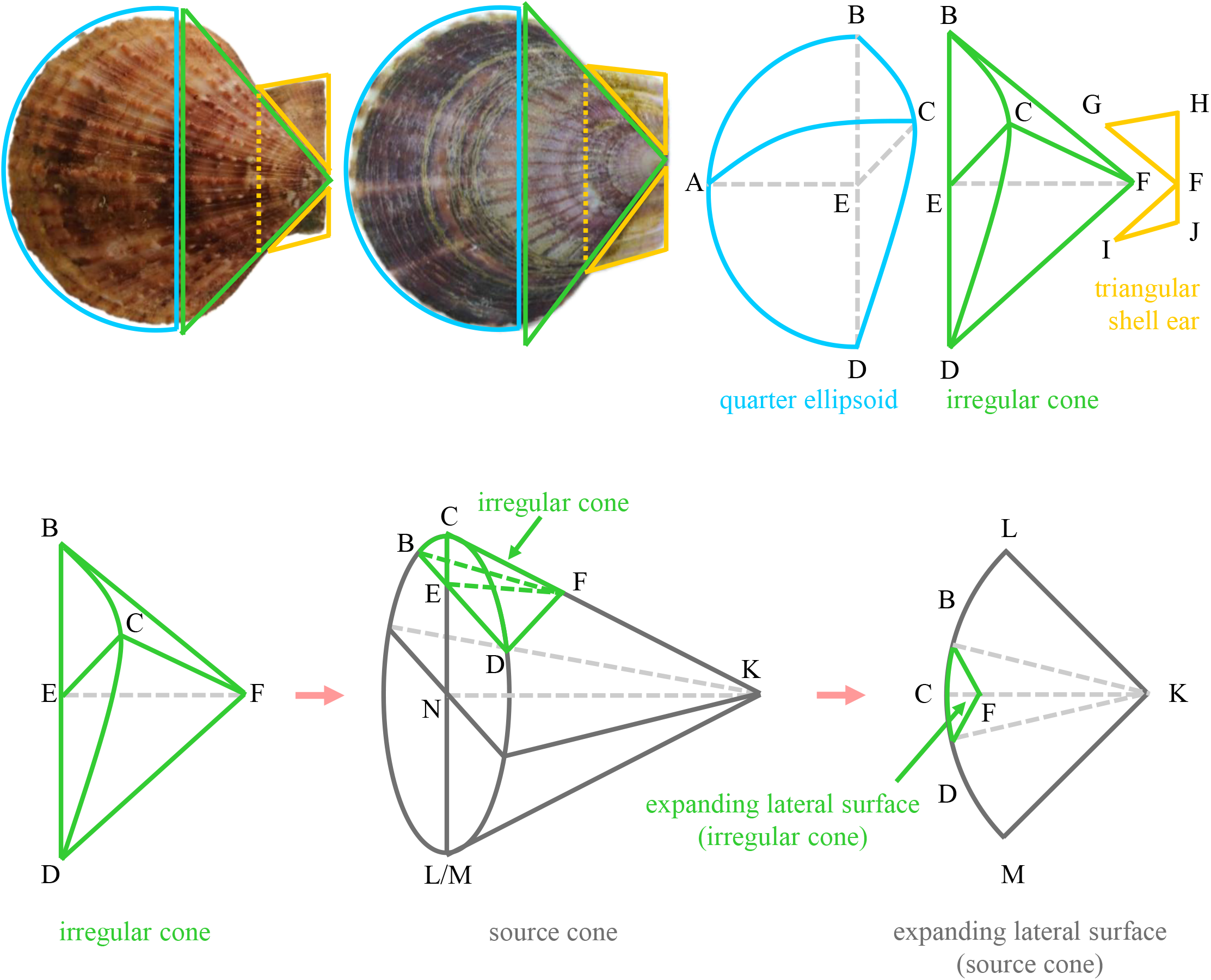
Demonstration of shell surface area calculation.

where *SL* and *SH* are shell length and shell height, respectively, *SW*_1_ and *SW*_2_ are widths for corresponding shells, *H*_*sb*1_ and *H*_*sb*2_ are heights of shell bottoms for corresponding shells, and *L*_*sb*11_ and *L*_*sb*12_ are lengths of upper bases of shell bottoms for corresponding shells.

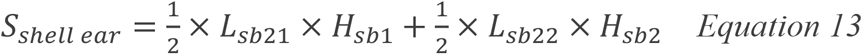

where *H*_*sb*1_ and *H*_*sb*2_ are heights of shell bottoms for corresponding shells, and *L*_*sb*21_ and *L*_*sb*22_ are lengths of lower bases for corresponding shells.

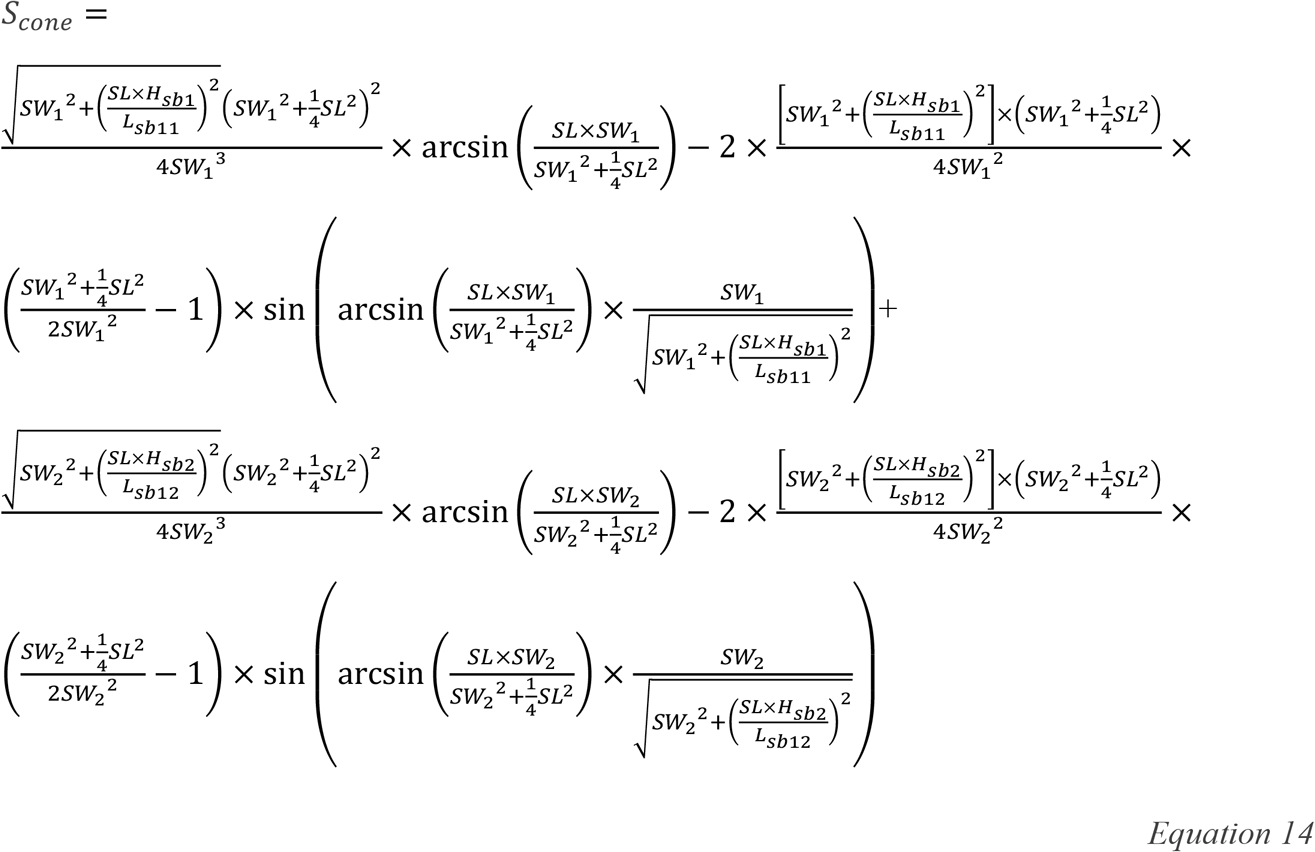

We fitted wet weight and shell surface area, as independent variables, and soft tissue weight, as a dependent variable, of samples in the training set to multiple linear regression (Equation 11) to obtain values of coefficients, a, b, and c, using OLS regression, as implemented in R 4.3.0 (R Core Team, 2023), and ridge regression, as implemented in SPSSPRO. With a, b, and c estimated from the training set, we calculated soft tissue weight for testing sets using Equation 11 and named this multiple linear regression-based estimation as MLR-*S*_*shell*_-OLS and MLR-*S*_*shell*_-RR methods for estimating soft tissue weight using OLS and ridge regression, respectively.

We further fitted log-transformed wet weight, log_10_ *W*_*wet*_, and log-transformed shell surface area, log_10_ *S*_*shell*_, as independent variables, and log-transformed soft tissue weight, log_10_ *W*_*soft*_, as a dependent variable, of samples in the training set to multiple linear regression (Equation 15) to obtain values of coefficients, a, b, and c, using OLS regression, as implemented in R 4.3.0 (R Core Team, 2023), and ridge regression, as implemented in SPSSPRO. With a, b, and c estimated from the training set, we calculated soft tissue weight for testing sets using Equation 15 and named this multiple linear regression-based estimation as MLR-log_10_ *S*_*shell*_-OLS and MLR-log_10_ *S*_*shell*_-RR methods for estimating soft tissue weight using OLS and ridge regression, respectively.

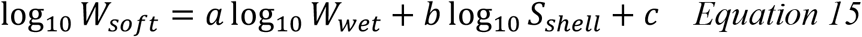

where a, b, and c are constants.

Some measurements, such as *L*_*sb*11_, *L*_*sb*12_, *L*_*sb*21_, *L*_*sb*22_, *H*_*sb*1_, *H*_*sb*2_, *SW*_1_, and *SW*_2_, that are needed in calculating shell surface area can hardly be made in practice on a large scale. In contrast, shell length, shell width, and shell height can be easily measured, so we fitted multiple linear regression to shell length, shell width, and shell height as independent variables and the estimated shell surface area as a dependent variable (Equation 16).

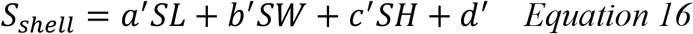

where *a*^′^, *b*^′^, *c*^′^, and *d*^′^ are constants.

By substituting shell surface area, *S*_*shell*_, in Equation 11 with *a*^′^*SL* + *b*^′^*SW* + *c*^′^*SH* + *d*^′^ in Equation 16, we can estimate soft tissue weight with shell length, shell width, and shell height directly (Equation 17). We fitted wet weight, shell length, shell width, and shell height, as independent variables, and soft tissue weight, as a dependent variable, of samples in the training set to multiple linear regression (Equation 17) to obtain values of coefficients, *a, a*^′^*b, b*^′^*b, bc*^′^, and *bd*^′^ + *c*, using OLS regression, as implemented in R 4.3.0 (R Core Team, 2023), and ridge regression, as implemented in SPSSPRO. With *a, a*^′^*b, b*^′^*b, bc*^′^, and *bd*^′^ + *c* estimated from the training set, we calculated soft tissue weight for testing sets using Equation 17 and named this multiple linear regression-based estimation as approximate MLR-*S*_*shell*_-OLS and approximate MLR-*S*_*shell*_-RR methods for estimating soft tissue weight using OLS and ridge regression, respectively.

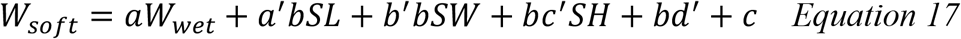

where *a, a*^′^*b, b*^′^*b, bc*^′^, and *bd*^′^ + *c* are constants.

We further took base 10 logarithmic transformations for all variables in Equation 17 to fit log-transformed wet weight, shell length, shell width, and shell height, as independent variables, and log-transformed soft tissue weight, as a dependent variable, of samples in the training set to multiple linear regression (Equation 18) to obtain values of coefficients, *a, a*^′^*b, b*^′^*b, bc*^′^, and *bd*^′^ + *c*, using OLS regression, as implemented in R 4.3.0 (R Core Team, 2023), and ridge regression, as implemented in SPSSPRO. With *a, a*^′^*b, b*^′^*b, bc*^′^, and *bd*^′^ + *c* estimated from the training set, we calculated soft tissue weight for testing sets using Equation 18 and named this multiple linear regression-based estimation as approximate MLR-log_10_ *S*_*shell*_-OLS and approximate MLR-log_10_ *S*_*shell*_-RR methods for estimating soft tissue weight using OLS and ridge regression, respectively.

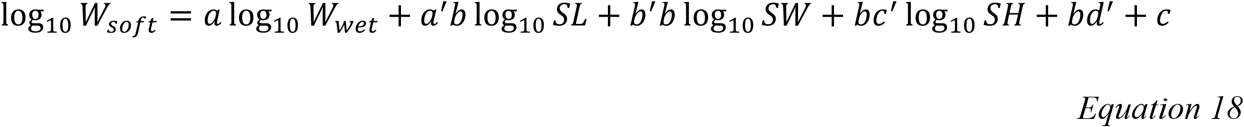

### 2.4 Accuracy of soft tissue weight estimation

To assess the accuracy of shell dimension-, wet weight-, and shell surface area-based in vivo assessment of soft tissue weight, we first constructed a simple linear regression model of estimated soft tissue weight as a dependent variable against observed soft tissue weight as an independent variable and calculated Pearson and Spearman correlation coefficients in R 4.3.0 (R Core Team, 2023) to evaluate how soft tissue weight estimated by different methods correlate with the observed values. We further tested whether observed and estimated soft tissue weight are significantly different using paired two-sample t-test and paired samples Wilcoxon signed rank test given the violation of normality (see 3.1 below for details in normality tests) in R 4.3.0 (R Core Team, 2023). Finally, we calculated the mean squared error (MSE), the average squared difference between estimated and observed values, and Cohen’s *d*, an assessment of effect size, of different models to assess the size of differences.

## 3 Results

### 3.1 Summary of shell dimensions and wet weight of scallops

In Zhikong scallop, population *Cf1* have significantly larger shell length (*Cf1* vs. *Cf2* mean shell length ± S.D.: 67.12 ± 6.01 vs. 54.62 ± 4.53 mm), shell width (*Cf1* vs. *Cf2* mean shell width ± S.D.: 25.46 ± 2.16 vs. 16.14 ± 1.54 mm), shell height (*Cf1* vs. *Cf2* mean shell height ± S.D.: 72.52 ± 5.84 vs. 50.44 ± 4.52 mm), and wet weight (*Cf1* vs. *Cf2* mean wet weight ± S.D.: 54.77 ± 11.64 vs. 19.99 ± 4.37 g) than population *Cf2* on average (*P* < 0.0001; Fig. 3, Table S2, Table S3). In yesso scallop, populations *Hg* and *Zr* have similar shell length (*P* = 0.7927; *Hg* vs. *Zr* mean shell length ± S.D.: 97.96 ± 12.71 vs. 95.85 ± 6.49 mm) and shell height (*P* = 0.1810; *Hg* vs. *Zr* mean shell height ± S.D.: 99.70 ± 11.75 vs. 96.74 ± 5.93 mm; Fig. 3, Table S3), but significantly differ in shell width (*P* = 0.0016; *Hg* vs. *Zr* mean shell width ± S.D.: 28.15 ± 3.44 vs. 26.76 ± 2.09 mm) and wet weight (*P* = 0.0007; *Hg* vs. *Zr* mean wet weight ± S.D.: 126.90 ± 44.09 vs. 108.48 ± 21.26 g; Fig. 3, Table S3). Zhikong and yesso scallops all exhibit substantial variation in shell dimensions and wet weight among individuals within each population (Fig. 3, Table S2). Only three (i.e., shell width in *Cf1* and *Zr*, wet weight in *Cf2*) and seven (i.e., shell length in *Cf1* and *Zr*, shell width in *Cf1, Cf2*, and *Zr*, wet weight in *Cf1* and *Cf2*) traits of shell dimensions and wet weight are normally distributed at *α* = 0.05 and *α* = 0.01, respectively (Table S2).

**Figure 3.**
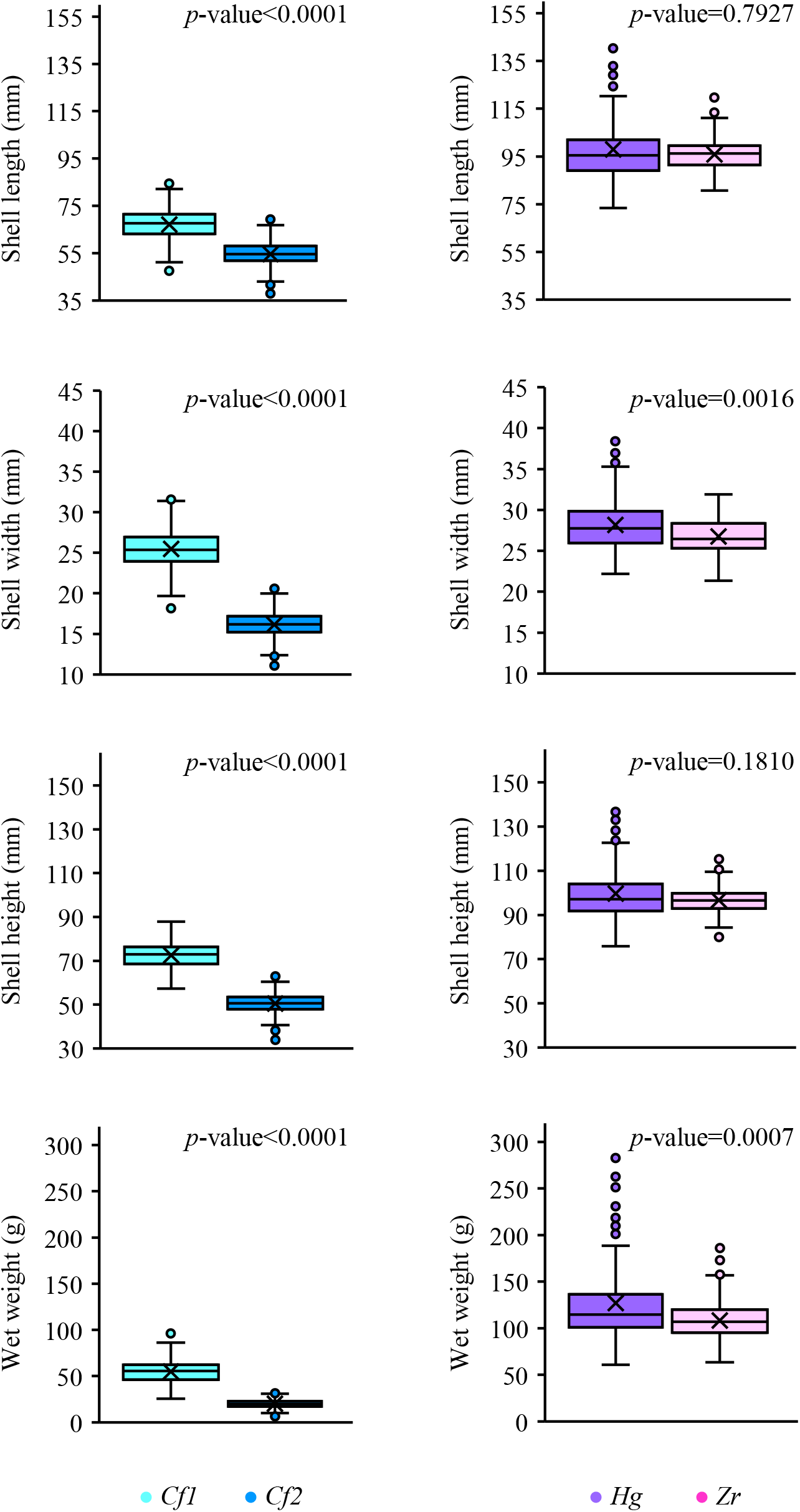
Boxplots of shell length, shell width, shell height, and wet weight of two Zhikong (*Cf1, Cf2*) and two yesso (*Hg, Zr*) scallop populations. In the Boxplots, the middle line of the box represents the median, symbol “×” in the box represents the mean, the bottom line of the box represents the first quartile, and the top line of the box represents the third quartile. The distance between the first and the third quartiles is the interquartile range (IQR). Solid points represent outliers that exceed 1.5 times of IQR below the first or above the third quartile. Whiskers (vertical lines) extend from the bottom and top lines of the box to the minimum and maximum value of the dataset excluding outliers. *P* value of Mann-Whitney U test indicates whether two populations within the same species differ in shell length, shell width, shell height, and wet weight.

### 3.2 Accuracy of in vivo assessment of soft tissue weight of scallops

#### 3.2.1 Pearson correlation coefficients between observed and estimated soft tissue weight

Pearson correlation coefficients (*r*) between soft tissue weight observed and estimated by 13 models constructed using OLS and ridge regression vary from 0.677 to 0.975 for 72 comparisons (i.e., 2 training sets×2 testing sets×(13 models of OLS regression+5 models of ridge regression)=72) in Zhikong scallop and range from 0.516 to 0.972 for 72 comparisons in yesso scallop (Fig. 4, Table S4). Over 8%, 2%, 33%, and 55% in Zhikong scallop and over 11%, 1%, 11%, and 65% in yesso scallop of 72 comparisons have Pearson correlation coefficients equal to or greater than 0.6, 0.7, 0.8, and 0.9, respectively (Fig. 4, Table S4). Pearson and Spearman correlation coefficients of all 144 comparisons in both Zhikong and yesso scallops are highly correlated (*r*=0.983; Table S4).

**Figure 4.**
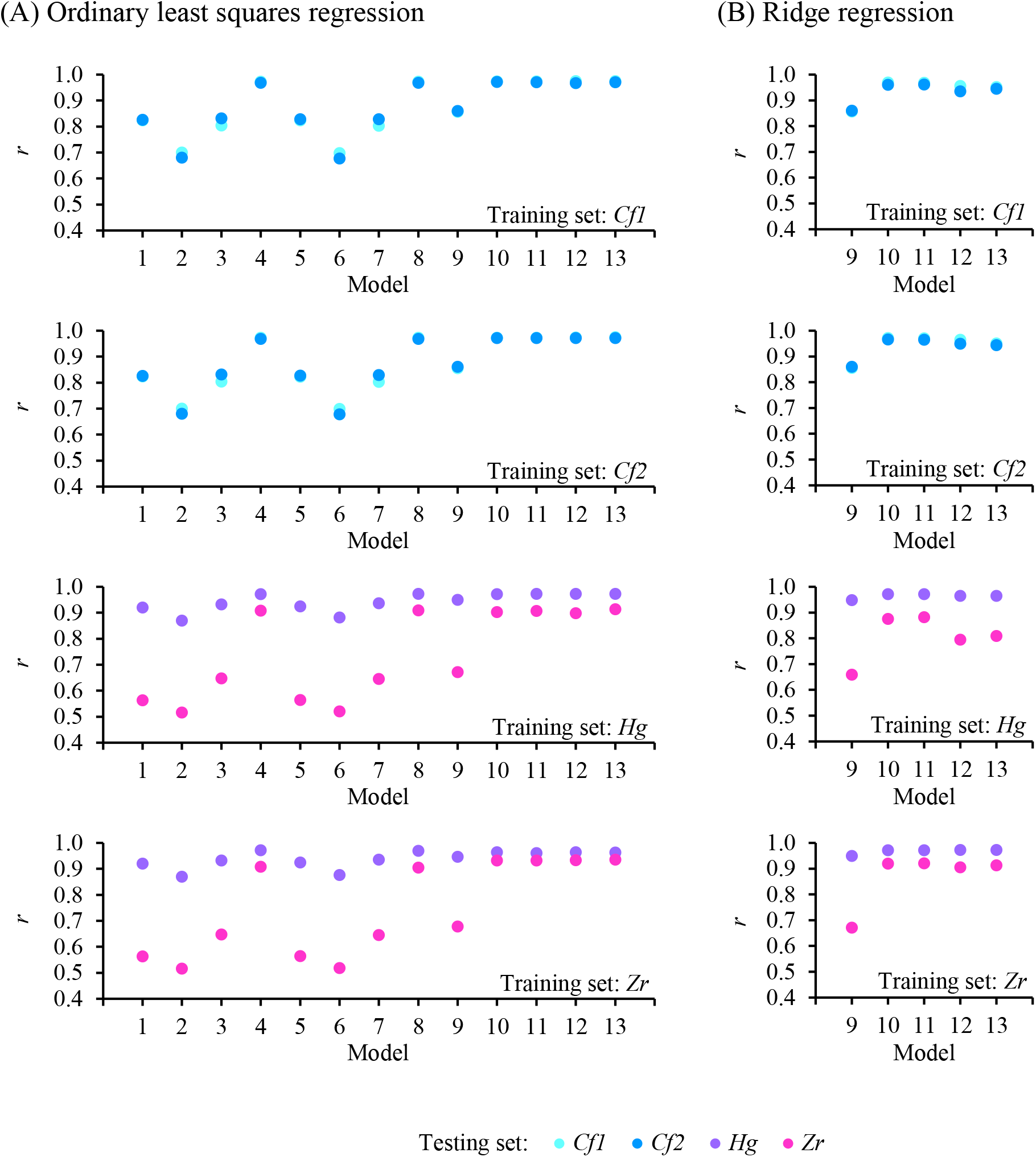
Pearson correlation coefficients (*r*) between soft tissue weight observed and estimated by 13 models constructed using OLS (A) and ridge (B) regression for different training and testing sets. Model 1: SLR-*SL*, Model 2: SLR-*SW*, Model 3: SLR-*SH*, Model 4: SLR-*W*_*wet*_, Model 5: PF-*SL*, Model 6: PF-*SW*, Model 7: PF-*SH*, Model 8: PF-*W*_*wet*_, Model 9: MLR-*SLWH*-OLS or MLR-*SLWH*-RR; Model 10: MLR-*S*_*shell*_-OLS or MLR-*S*_*shell*_-RR; Model 11: MLR-log_10_ *S*_*shell*_-OLS or MLR-log_10_ *S*_*shell*_-RR; Model 12: Approximate MLR-*S*_*shell*_-OLS or Approximate MLR-*S*_*shell*_-RR; Model 13: Approximate MLR-log_10_ *S*_*shell*_-OLS or Approximate MLR-log_10_ *S*_*shell*_-RR. See Material and methods for explanations for models and populations.

For models based on simple linear regression and power function with a single trait (e.g., SLR-*SL*, SLR-*SW*, SLR-*SH*, SLR-*W*_*wet*_, PF-*SL*, PF-*SW*, PF-*SH*, PF-*W*_*wet*_), wet weight-based estimates (*r*: 0.969-0.973 and 0.904-0.972 for Zhikong and yesso scallops, respectively; Fig. 4, Table S4) are more closely correlated with observed soft tissue weight than those predicted based on a single shell dimension (*r*: 0.677-0.831 and 0.516-0.936 for Zhikong and yesso scallops, respectively; Fig. 4, Table S4). For multiple linear regression-based methods, shell surface area-based models incorporating both shell surface area and wet weight (e.g., MLR-*S*_*shell*_-OLS, MLR-*S*_*shell*_-RR, MLR-log_10_ *S*_*shell*_-OLS, MLR-log_10_ *S*_*shell*_-RR, approximate MLR-*S*_*shell*_-OLS, approximate MLR-*S*_*shell*_-RR, approximate MLR-log_10_ *S*_*shell*_-OLS, approximate MLR-log_10_ *S*_*shell*_-RR; *r*: 0.935-0.975 and 0.795-0.972 for Zhikong and yesso scallops, respectively) outperform those built on shell length, shell width, and shell height (e.g., MLR-*SLWH*-OLS, MLR-*SLWH*-RR; *r*: 0.856-0.861 and 0.659-0.949 for Zhikong and yesso scallops, respectively) in estimating soft tissue weight (Fig. 4, Table S4). For shell surface area-based models, OLS (*r*: 0.967-0.975 and 0.898-0.972 for Zhikong and yesso scallops, respectively) and ridge (*r*: 0.935-0.972 and 0.795-0.972 for Zhikong and yesso scallops, respectively) regression-based methods perform nearly equivalently well, except for a slightly poorer estimation of *Zr* soft tissue weight by approximate MLR-*S*_*shell*_-RR and approximate MLR-log_10_ *S*_*shell*_-RR with *Hg* as the training set (*r*: 0.795 and 0.809; Fig. 4, Table S4). Overall, correlation coefficients of observed soft tissue weight with estimates by wet weight-involving models, ranging from 0.935 to 0.975 in Zhikong scallop and from 0.795 to 0.972 in yesso scallops, are higher than those by models constructed using only shell dimensions (*r*: 0.677-0.861 and 0.516-0.949 in Zhikong and yesso scallops, respectively; Fig. 4, Table S4).

#### 3.2.2 Paired samples Wilcoxon signed rank tests for soft tissue weight

To further test how accurate the estimation is, we conducted paired samples Wilcoxon signed rank tests for each pair of observed and predicted soft tissue weight in 72 comparisons for Zhikong scallop and 72 comparisons for yesso scallops given the violation of normality. Soft tissue weight of a population can be accurately predicted using models constructed with the same population as the training set (*P* > 0.05) except for the estimation of soft tissue weight in *Zr* by the approximate MLR-log_10_ *S*_*shell*_-RR model trained by the same population, in which observed and estimated soft tissue weight are not significantly different at *α* = 0.01 but barely different at *α* = 0.05 (P = 0.041; Table S4). In cases of estimating soft tissue weight for one testing set with a different training set, 2 and 14 out of 36 estimations are not significantly different than observed soft tissue weight at *α* = 0.05 in Zhikong and yesso scallops, respectively (Table S4). Given that the strength of associations and the exact differences between observed and estimated soft tissue weight are both critical metrics for evaluating the prediction accuracy of models, we identified candidate models as those satisfying the following two criteria: (1) correlation coefficients between observed and estimated soft tissue weight above 0.8, and (2) *P* values of paired samples Wilcoxon signed rank tests for observed and estimated soft tissue weight greater than 0.05 (Table 1). MLR-*log*_10_*S*_*shell*_-RR is the only candidate model for Zhikong scallop, which can accurately predict soft tissue weight for one population using the other as the training set. For yesso scallop, only models MLR-*log*_10_*S*_*shell*_-RR and approximate MLR-*log*_10_*S*_*shell*_-RR works for estimating soft tissue weight of *Hg* with *Zr* as the training set and vice versa, while none of the other models can generate reliable assessments when training and testing sets are switched. Therefore, MLR-*log*_10_*S*_*shell*_-RR is the optimal model in comparison to all others tested in this study as it accommodates training sets from both Zhikong and yesso scallops, generates reliable estimates for both species, and performs equivalently well when training and testing sets are switched (Table 1).

**Table 1.** Pearson correlation coefficients (*r*) and *P* values of paired samples Wilcoxon signed rank test for differences between soft tissue weight observed and estimated by different models for different training and testing sets. See Material and methods for explanations for models and populations.

| Model | Training set | Testing set | $r$ | $P$ value |
| --- | --- | --- | --- | --- |
| MLR-$\log_{10} S_{shell}$-RR | *Cf1* | *Cf2* | 0.9616 | 0.3649 |
| MLR-$\log_{10} S_{shell}$-RR | *Cf2* | *Cf1* | 0.9701 | 0.3392 |
| SLR-$W_{wet}$ | *Hg* | *Zr* | 0.9082 | 0.1853 |
| MLR-$S_{shell}$-OLS | *Hg* | *Zr* | 0.9030 | 0.2936 |
| MLR-$S_{shell}$-RR | *Hg* | *Zr* | 0.8759 | 0.9460 |
| MLR-$\log_{10} S_{shell}$-RR | *Hg* | *Zr* | 0.8825 | 0.3807 |
| MLR-$\log_{10} S_{shell}$-RR | *Zr* | *Hg* | 0.9711 | 0.0750 |
| Approximate MLR-$S_{shell}$-OLS | *Hg* | *Zr* | 0.8983 | 0.3704 |
| Approximate MLR-$S_{shell}$-RR | *Zr* | *Hg* | 0.9720 | 0.1815 |
| Approximate MLR-$\log_{10} S_{shell}$-RR | *Hg* | *Zr* | 0.8088 | 0.8404 |
| Approximate MLR-$\log_{10} S_{shell}$-RR | *Zr* | *Hg* | 0.9721 | 0.6611 |

#### 3.2.3 Mean squared errors and Cohen’s *d*

To verify the reliability and robustness of MLR-*log*_10_*S*_*shell*_-RR model, we calculated mean squared errors (MSE) and Cohen’s *d* for soft tissue weight observed and estimated by 13 models constructed using OLS and ridge regression for the four training set-testing set combinations for which model MLR-*log*_10_*S*_*shell*_-RR works well (i.e., training set: *Cf1*, testing set: *Cf2*; training set: *Cf2*, testing set: *Cf1*; training set: *Hg*, testing set: *Zr*; training set: *Zr*, testing set: *Hg*). MSE of model MLR-*log*_10_*S*_*shell*_-RR is nearly the lowest in comparison to that of all other models except for the *Hg*-*Zr* combination, in which MSE of MLR-*log*_10_*S*_*shell*_-RR model is slightly higher than that of models involving wet weight but is about two times lower than that of models based on only shell dimensions (Fig. 5, Table S4). Cohen’s *d* of model MLR-*log*_10_*S*_*shell*_-RR is nearly zero in all combinations (Fig 5, Table S4). A scrutiny of plots of ranked observed soft tissue weight against its estimates by wet weight-involving models, SLR-*W*_*wet*_, PF-*W*_*wet*_, and MLR-*log*_10_*S*_*shell*_-RR, for the four training set-testing set combinations of Zhikong and yesso scallops above reveals that SLR-*W*_*wet*_ and PF-*W*_*wet*_ tend to systematically overestimate (i.e., PF-*W*_*wet*_ for *Cf2* as training set and *Cf1* as testing set, SLR-*W*_*wet*_ and PF-*W*_*wet*_ for *Zr* as training set and *Hg* as testing set) or underestimate (i.e., SLR-*W*_*wet*_ and PF-*W*_*wet*_ for *Cf1* as training set and *Cf2* as testing set) soft tissue weight in comparison to model MLR-*log*_10_*S*_*shell*_-RR (Fig. 6). Such over- or under-estimations by various models are pervasive despite the high correlation between observed and estimated soft tissue weight for both Zhikong and yesso scallops (Fig. S1).

**Figure 5.**
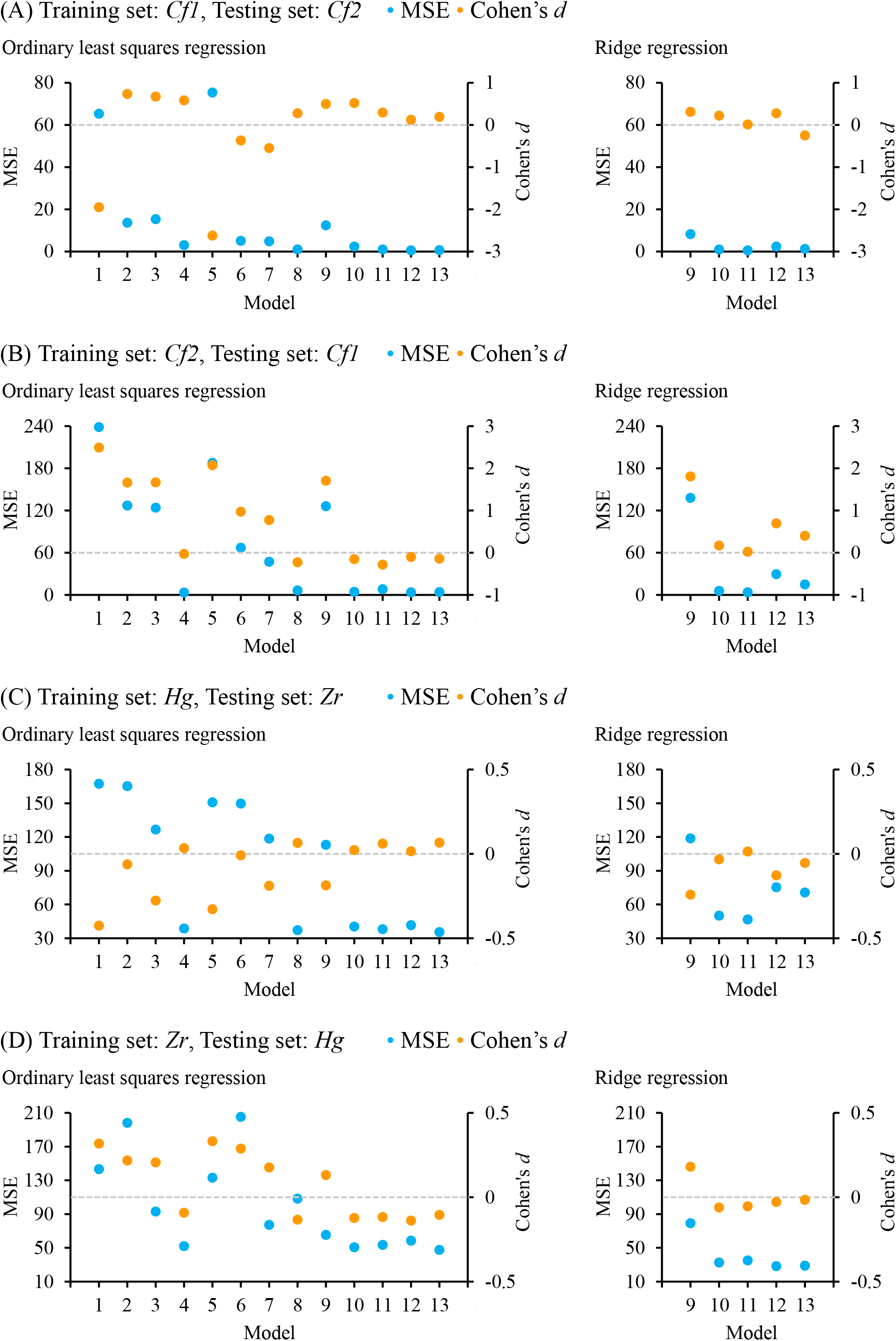
Mean squared errors (MSE) and Cohen’s *d* for 13 models constructed using OLS and ridge regression for four training set-testing set combinations. (A) *Cf1* as training set and *Cf2* as testing set, (B) *Cf2* as training set and *Cf1* as testing set, (C) *Hg* as training set and *Zr* as testing set, (D) *Zr* as training set and *Hg* as testing set. Dashed, horizontal, grey line indicates Cohen’s *d* of 0. Model 1: SLR-*SL*, Model 2: SLR-*SW*, Model 3: SLR-*SH*, Model 4: SLR-*W*_*wet*_, Model 5: PF-*SL*, Model 6: PF-*SW*, Model 7: PF-*SH*, Model 8: PF-*W*_*wet*_, Model 9: MLR-*SLWH*-OLS or MLR-*SLWH*-RR; Model 10: MLR-*S*_*shell*_-OLS or MLR-*S*_*shell*_-RR; Model 11: MLR-log_10_ *S*_*shell*_-OLS or MLR-log_10_ *S*_*shell*_-RR; Model 12: Approximate MLR-*S*_*shell*_-OLS or Approximate MLR-*S*_*shell*_-RR; Model 13: Approximate MLR-log_10_ *S*_*shell*_-OLS or Approximate MLR-log_10_ *S*_*shell*_-RR. See Material and methods for explanations for models and populations.

**Figure 6.**
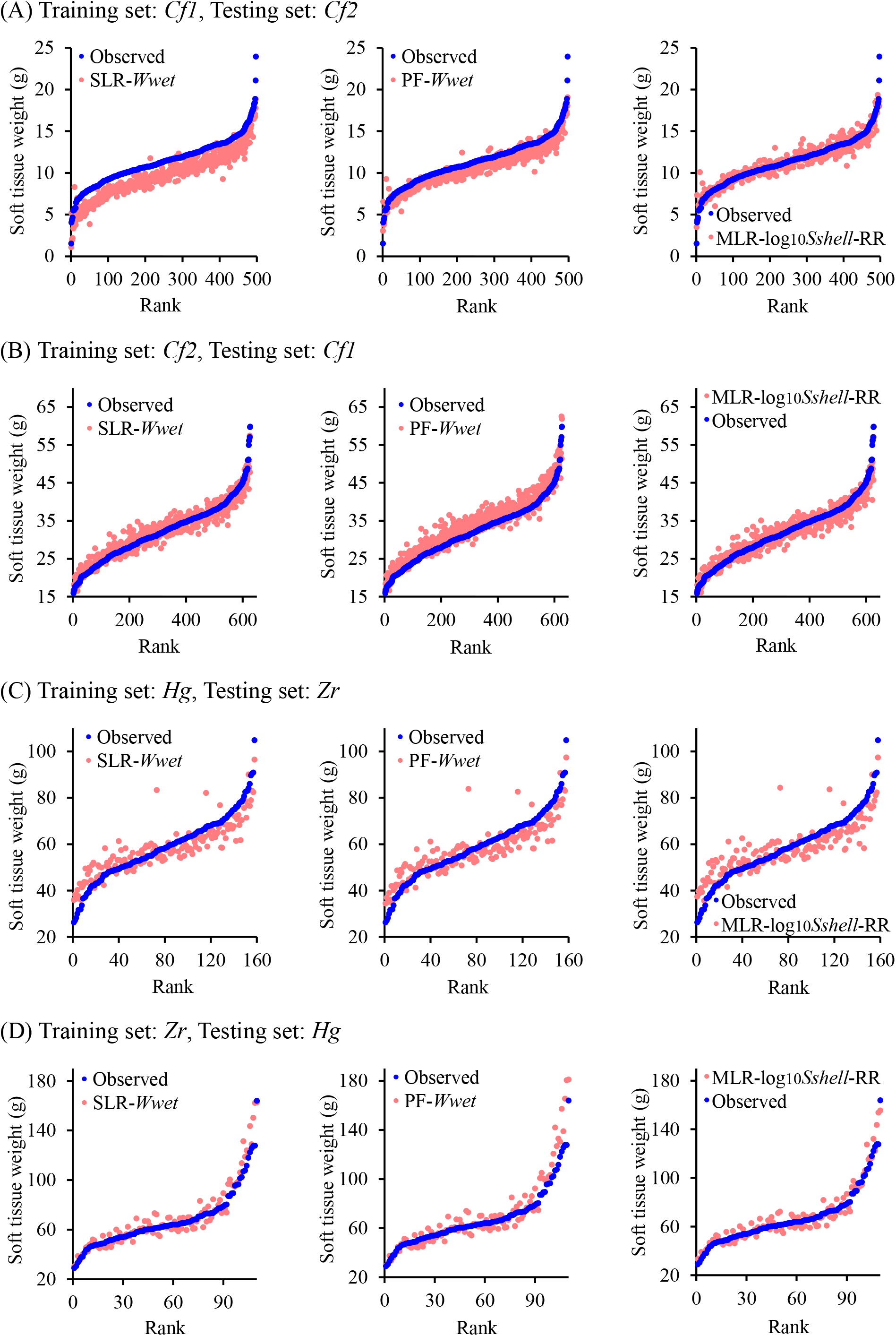
Scatterplots of ranked observed soft tissue weight against corresponding estimates by models SLR-*W*_*wet*_, PF-*W*_*wet*_, and MLR-log_10_ *S*_*shell*_-RR for four training set-testing set combinations. (A) *Cf1* as training set and *Cf2* as testing set, (B) *Cf2* as training set and *Cf1* as testing set, (C) *Hg* as training set and *Zr* as testing set, (D) *Zr* as training set and *Hg* as testing set. See Material and methods for explanations for models and populations.

### 3.3 Soft tissue weight estimation with subsets of full training sets

To test whether the optimal model, MLR-*log*_10_*S*_*shell*_-RR, still performs well with a smaller training set, we take subsets of *Cf1* (n=50, 100, 150, 200, 250, 300) and *Cf2* (n=50, 100, 150, 200) as training sets to estimate soft tissue weight for *Cf2* and *Cf1*, as testing sets, respectively. Even though smaller subsets are used for model construction, observed soft tissue weight and its estimates by MLR-*log*_10_*S*_*shell*_-RR are still highly correlated in both *Cf1* and *Cf2*, with correlation coefficients of 0.956-0.966 for estimating *Cf2* with subsets of *Cf1* as training sets and 0.968-0.973 for estimating *Cf1* with subsets of *Cf2* as training sets (Fig. 7). Variation in Pearson correlation coefficients of three replicated estimations with training sets of 50 samples is greater than that with larger training sets (Fig. 7). Observed and estimated *Cf2* soft tissue weights are not significantly different at *α* = 0.05 in one replicated estimation using *Cf1* of 50 samples as the training set and in two replicated estimations using *Cf1* of 250 and 300 samples as training sets (Fig. 7). In contrast, all estimations of *Cf1* soft tissue weight are significantly different than actual values at *α* = 0.05 when subsets of *Cf2* are used as training sets (Fig. 7). Systematic over- or under-estimations with smaller training sets may account for the disparity between observed and predicted soft tissue weight (Fig. S2).

**Figure 7.**
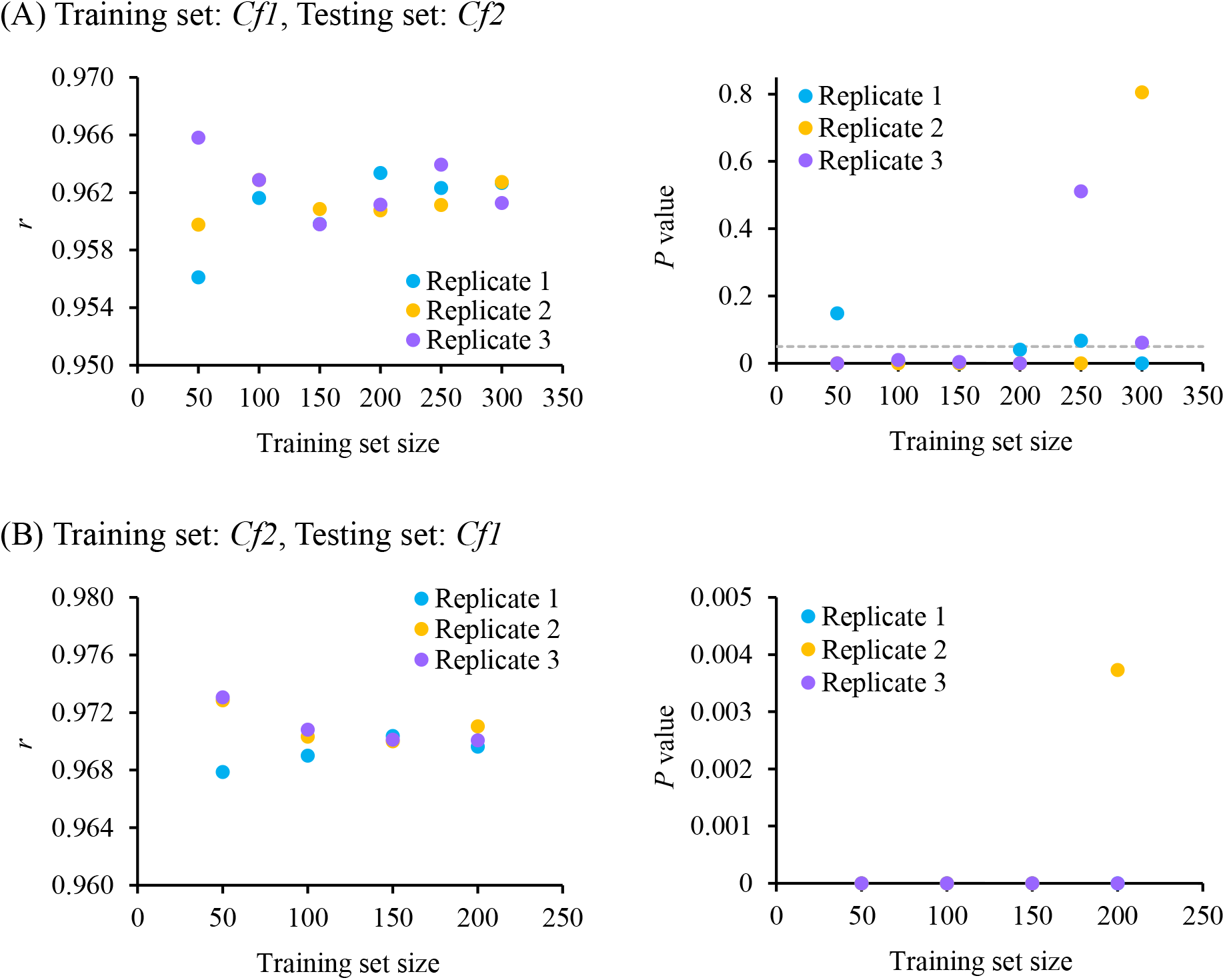
Pearson correlation coefficients (*r*) and *P* values of paired samples Wilcoxon signed rank test for differences between soft tissue weight observed and estimated by model MLR-log_10_ *S*_*shell*_-RR constructed using three replicated subsets of *Cf1* (A; n=50, 100, 150, 200, 250, 300) and *Cf2* (B; n=50, 100, 150, 200) as training sets for estimating soft tissue weight for the other population. See Material and methods for explanations for models and populations. Dashed, horizontal, grey line indicates *P* value of 0.05.

## 4 Discussion

Rapid, reliable, in vivo assessment of commercially important traits is critical for accelerating the progress of genetic improvements to enhance the efficiency of aquaculture production. However, traditional dissection- or imaging-based phenotyping methods are unable to make an accurate but nondestructive assessment of target traits, such as soft tissue weight, in bivalves. To resolve this issue, we decipher the complex relationships among all types of shell dimensions and weights, which have mostly been investigated in bivalves from an ecological perspective, and take advantage of such relationships to estimate soft tissue weight directly. Specifically, we develop a novel, robust, shell surface area-based method for a rapid and noninvasive assessment of soft tissue weight, which works for two predominant aquaculture species in China, Zhikong and yesso scallops, and similar bivalves with bilaterally symmetrical shells. In comparison to classical models, the shell surface area-based method accommodates collinearity among a collection of phenotypic traits, including various shell dimensions and weights, using ridge regression and eliminates systematic biases in soft tissue weight estimation.

Correlation coefficients of observed soft tissue weight with estimates by models based on only shell dimensions varies in a wider range in comparison to that with estimates by wet weight-involving methods, suggesting that wet weight is a key proxy for soft tissue weight and shell dimensions alone may not provide sufficient information for estimation. The inaccurate estimation by models based on only shell dimensions could be due to the large variation in adductor muscle size and soft tissue weight across scallops with similar sizes (Zhao et al., 2021). That wet weight-based estimates are always highly correlated with observed values does not necessarily mean all wet weight-involving models are reliable, as correlation coefficients only tells the strength of associations between observed and predicted values but cannot assess the estimation accuracy. Models incorporating wet weight, including SLR-*W*_*wet*_, PF-*W*_*wet*_, and shell surface area-based models fitted by both OLS and ridge regression, tend to over- or under-estimate soft tissue weight in 45% and 27% of cases in Zhikong and yesso scallops, respectively, even though the observed soft tissue weight is strongly associated with its estimates. Such pervasive, systematic biases may arise from the differential relationships among shell dimension- and weight-related traits across populations. Despite such complex relationships, model MLR-*log*_10_*S*_*shell*_-RR can accurately assess soft tissue weight for both Zhikong and yesso scallops, suggesting it as a robust model suitable for estimation in multiple species.

Model MLR-*log*_10_*S*_*shell*_-RR is superior in two aspects: first, model fitting can be conducted using training sets from both Zhikong and yesso scallops, with which an reliable estimation can be generated for both species; second, pairwise, reciprocal estimations are possible with this model, and specifically, *Cf1* and *Hg* can serve as training sets for estimating *Cf2* and *Zr*, respectively, and vice versa. Model MLR-*log*_10_*S*_*shell*_-RR estimates soft tissue weight by subtracting shell weight, which is the product of shell density, shell thickness, and shell surface area, from wet weight. As shell density and shell thickness are relatively constant across populations under the same environment, the relationship among shell surface area, wet weight, and soft tissue weight usually holds for multiple populations. Therefore, models constructed with one population can generate accurate estimation for another grown under similar environmental conditions.

Given the performance of model MLR-*log*_10_*S*_*shell*_-RR across training sets and populations, two factors appear to impact the estimation accuracy. One critical factor for soft tissue weight assessment is the size of training sets. The exact values of soft tissue weight of *Cf1* and *Cf2* cannot be accurately estimated by models constructed with subsets of complete *Cf2* and *Cf1* as training sets in most cases, suggesting that the size of training sets matters. While observed and estimated soft tissue weight are significantly different, they are still highly correlated, indicating that models based on smaller training sets are still able to make predictions but may over- or under-estimate actual values. In cases, for example parent line selection for breeding programs, where the rank, rather than the exact soft tissue weight, of candidates is important, estimations based on smaller training sets are still useful. It should also be noted that a training set of 50 samples from *Cf1* generates an accurate estimation for *Cf2*, implying that smaller training sets with an appropriate distribution may also work. Therefore, picking training sets of an appropriate size for estimations of different objectives is essential for model construction. The other factor that may affect the estimation accuracy is the genetic and environmental background. Shell morphology, shell density, and shell thickness may vary across genetically divergent populations and vastly different environments in some cases, restricting the potential of models based on one population or environment to apply to others. Thus, genetic and environmental background of training sets should be considered in model construction.

MLR-*log*_10_*S*_*shell*_-RR is a robust and reliable model for soft tissue weight assessment across populations not only for scallops but also for bivalves with similar morphology (e.g., bilaterally symmetrical shells), such as clams, mussels, and cockles. To facilitate the application of this shell surface area-based models, we construct approximate MLR-*log*_10_*S*_*shell*_-RR model, in which we substitute shell surface area with shell length, shell width, and shell height for which a large-scale measurement is feasible in practice. Such an approximate model works for yesso scallop but not for Zhikong scallop, suggesting that approximate model-based assessments are promising but further improvements for model training are needed. To enhance the performance of shell surface area-based models, we suggest developing more accurate, high-throughput, automatic methods for assessing morphometric traits, such as shell dimensions, to capture morphological characteristics at a finer scale, which will better inform model fitting. Meanwhile, a database incorporating training sets of various sizes that originate from different populations and species grown under multiple environments and their performance metrics, such as the correlation and difference between estimated and actual values, for all types of traits in testing sets of interest is essential for a broader application of this rapid, reliable, in vivo assessment method to boost bivalve phenotyping, breeding, and farming in the near future.

## Appendix A (See Fig. 2 for reference)

∵

*AF* = *SH, BD* = *SL*

*CE* = *SW*_1_ (*or CE* = *SW*_2_)

*GI* = *L*_*sb*11_ (*or GI* = *L*_*sb*12_)

*HJ* = *L*_*sb*21_ (*or HJ* = *L*_*sb*22_)

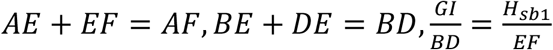

∵

1. AE and EF

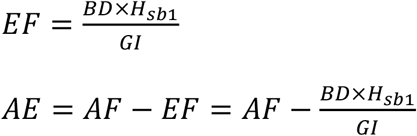
2. BE and DE

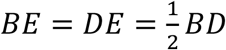

∵

1. CK, CN, and NK

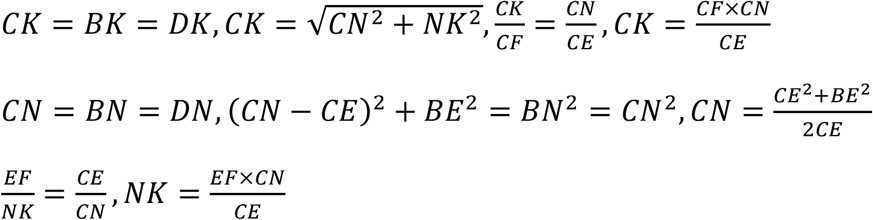
2. FK and CF

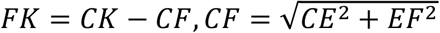
3. BF and DF

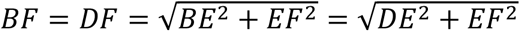
4. *θ*_∠*BKD*_ and *l*_*BCD*_

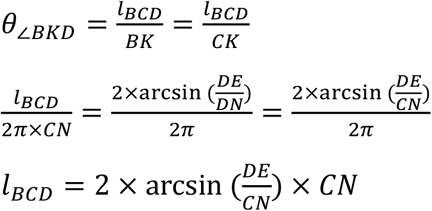
5. *θ*_∠*BKC*_

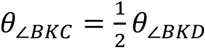
6. *S*_*sectorKBCD*_

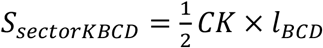
7. *S*_*triangleFBK*_ and *S*_*triangleFDK*_

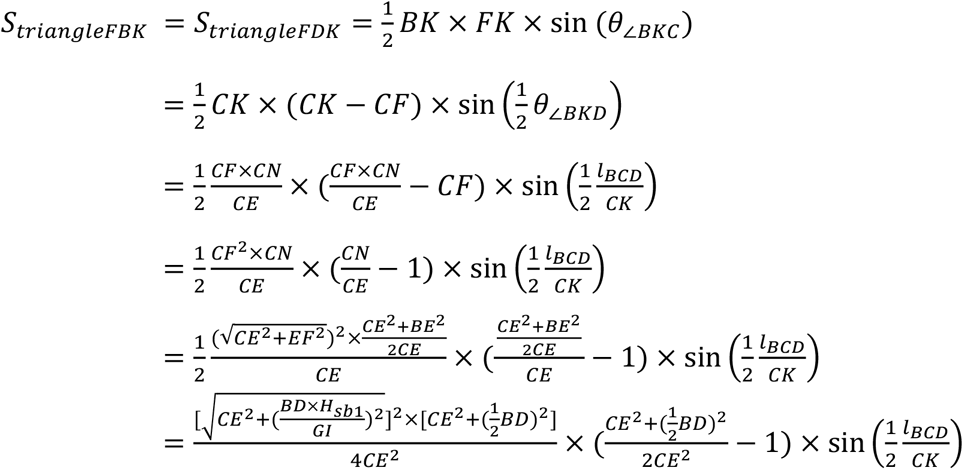

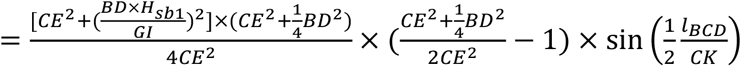
8. 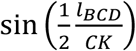

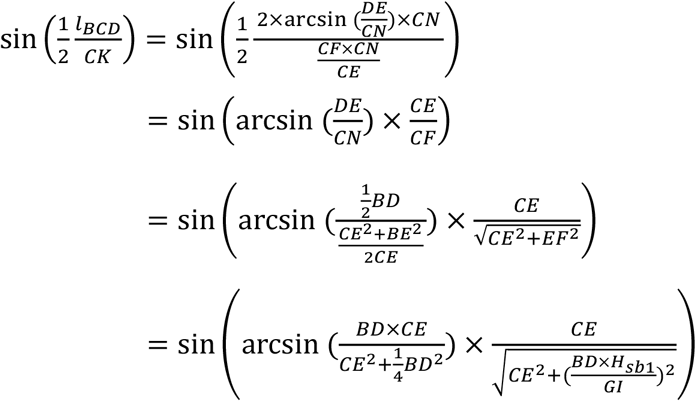
9. *S*_*sectorKBCD*_

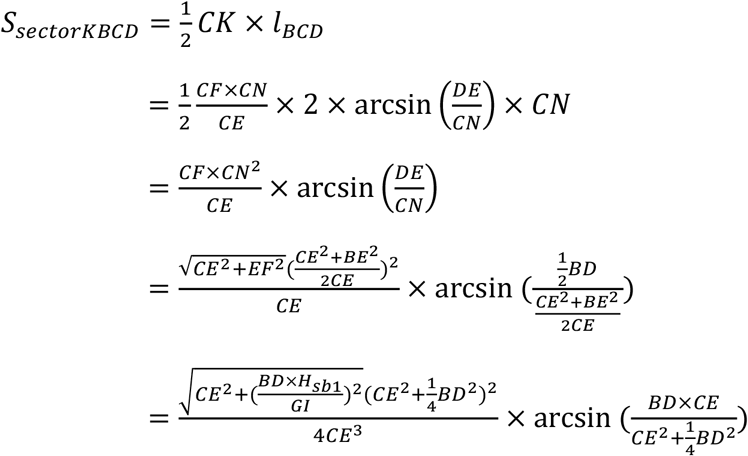
10. *S*_*sectorFBCD*_ and *S*_*cone*_

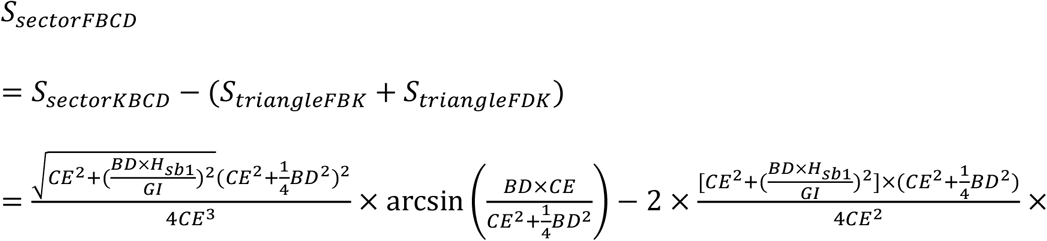

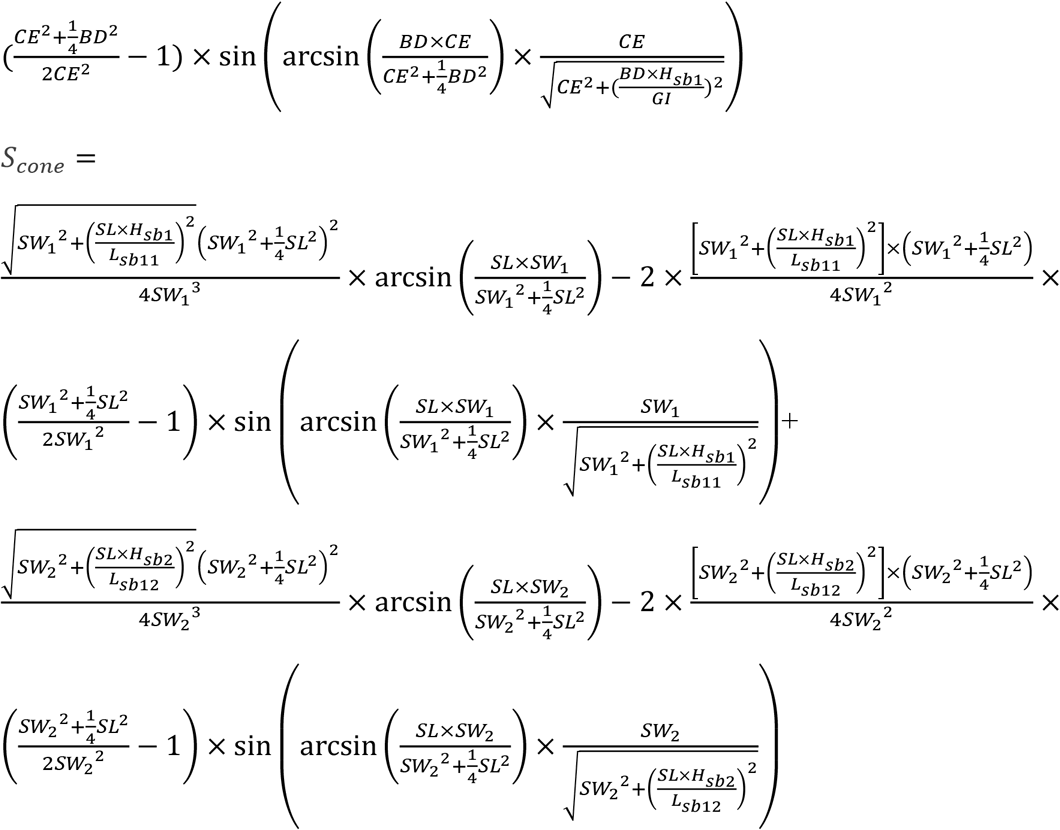

## Data statement

Raw data are available upon request.

## Author contributions: CRediT

Duan, S.: data curation, formal analysis, validation, writing-review and editing. Gao, Y.: data curation, formal analysis. Hu, X.: funding acquisition, writing-review and editing. Yin, X.: conceptualization, data curation, formal analysis, funding acquisition, investigation, methodology, project administration, supervision, writing-original draft, writing-review and editing.

## Acknowledgements

This work is supported by the High-performance Computing Platform of YZBSTCACC and the High-performance Biological Supercomputing Center at the Ocean University of China.

## References

Brand, E.V., Cisterna, M., Merino, G., Uribe, E., Palma-Rojas, C., Rosenblitt, M., Albornoz, J.L., 2009. Non-destructive method to study the internal anatomy of the Chilean scallop Argopecten purpuratus. Journal of Shellfish Research 28, 325–327. 10.2983/035.028.0214

Cameron, C.J., Cameron, I.F., Paterson, C.G., 1979. Contribution of organic shell matter to biomass estimates of unionid bivalves. Can. J. Zool. 57, 1666–1669. 10.1139/z79-217

Coughlan, N.E., Cunningham, E.M., Cuthbert, R.N., Joyce, P.W.S., Anastácio, P., Banha, F., Bonel, N., Bradbeer, S.J., Briski, E., Butitta, V.L., Čadková, Z., Dick, J.T.A., Douda, K., Eagling, L.E., Ferreira-Rodríguez, N., Hünicken, L.A., Johansson, M.L., Kregting, L., Labecka, A.M., Li, D., Liquin, F., Marescaux, J., Morris, T.J., Nowakowska, P., Ożgo, M., Paolucci, E.M., Peribáñez, M.A., Riccardi, N., Smith, E.R.C., Spear, M.J., Steffen, G.T., Tiemann, J.S., Urbanska, M., Van Doninck, K., Vastrade, M., Vong, G.Y.W., Wawrzyniak-Wydrowska, B., Xia, Z., Zeng, C., Zhan, A., Sylvester, F., 2021. Biometric conversion factors as a unifying platform for comparative assessment of invasive freshwater bivalves. Journal of Applied Ecology 58, 1945–1956. 10.1111/1365-2664.13941

Eklöf, J., Austin, Å., Bergström, U., Donadi, S., Eriksson, B.D.H.K., Hansen, J., Sundblad, G., 2017. Size matters: relationships between body size and body mass of common coastal, aquatic invertebrates in the Baltic Sea. PeerJ 5, e2906. 10.7717/peerj.2906

FAO, 2024. The state of world fisheries and aquaculture 2024. Blue transformation in action. Rome, FAO.

Golightly, C.G., Kosinski, R.J., 1981. Estimating the biomass of freshwater mussels (Bivalvia: Unionidae) from shell dimensions. Hydrobiologia 80, 263–267. 10.1007/BF00018366

Isom, B.G., 1971. The stepwise multiple regression method for selection of variables for predicting the shell weight of freshwater mussels. Malacological Review 4, 17–20.

Larson, J.H., Eckert, N.L., Bartsch, M.R., 2014. Intrinsic variability in shell and soft tissue growth of the freshwater mussel Lampsilis siliquoidea. PLOS ONE 9, e112252. 10.1371/journal.pone.0112252

McKinney, R.A., Glatt, S.M., Williams, S.R., 2004. Allometric length-weight relationships for benthic prey of aquatic wildlife in coastal marine habitats. Wildlife Biology 10, 241–249. 10.2981/wlb.2004.029

Michael Holliman, F., Davis, D., Bogan, A.E., Kwak, T.J., Gregory Cope, W., Levine, J.F., 2008. Magnetic resonance imaging of live freshwater mussels (Unionidae). Invertebrate Biology 127, 396–402. 10.1111/j.1744-7410.2008.00143.x

Molina, R., Hanlon, S., Savidge, T., Bogan, A., Levine, J., 2005. Buoyant weight technique: Application to freshwater bivalves. American Malacological Bulletin 20, 49–53.

Pouvreau, S., Rambeau, M., Cochard, J.C., Robert, R., 2006. Investigation of marine bivalve morphology by in vivo MR imaging: First anatomical results of a promising technique. Aquaculture 259, 415–423. 10.1016/j.aquaculture.2006.05.018

Powell, E.N., Mann, R., Ashton-Alcox, K.A., Kim, Y., Bushek, D., 2016. The allometry of oysters: spatial and temporal variation in the length-biomass relationships for Crassostrea virginica. Journal of the Marine Biological Association of the United Kingdom 96, 1127–1144. 10.1017/S0025315415000703

R Core Team, 2023. R: A language and environment for statistical computing. R Foundation for Statistical Computing, Vienna, Austria. https://www.R-project.org/.

Turra, A., Corte, G.N., Amaral, A.C.Z., Yokoyama, L.Q., Denadai, M.R., 2018. Non-linear curve adjustments widen biological interpretation of relative growth analyses of the clam Tivela mactroides (Bivalvia, Veneridae). PeerJ 6, e5070. 10.7717/peerj.5070

Wang, X., Liu, T., Liu, Y., Feng, P., 2014. An arithmetic index based on shell height, length, and width, for potential selection of soft-body wet weight in Pacific Oyster, Crassostrea gigas. The Israeli journal of aquaculture-Bamidgeh 66. 10.46989/001c.20671

Zhao, L., Lian, S., Ren, Q., Yang, Z., Guo, Z., Lou, J., Kong, X., Li, M., Bao, Z., Hu, X., 2021. In vivo and rapid assessment of scallop muscle trait. Aquaculture 530, 735817. 10.1016/j.aquaculture.2020.735817

